# Rapid evolution of learning and reproduction in natural populations of *Drosophila melanogaster*

**DOI:** 10.1101/288696

**Authors:** Emily L. Behrman, Tadeusz J. Kawecki, Paul Schmidt

**Author notes:** Corresponding Author: Emily L. Behrman Department of Biological Sciences, Dartmouth College, 78 College Street, Hanover, NH 03755. Authors contributed equally (Tadeusz J. Kawecki, Paul Schmidt).

## Abstract

Learning is a general mechanism of adaptive behavioural plasticity whose benefits and costs depend on the environment. Seasonal oscillations in temperate environments between winter and summer produce cyclical selection pressures that may drive rapid evolution of learning performance in multivoltine populations. To test this hypothesis, we investigated the evolutionary dynamics of learning ability over rapid seasonal timescales in a natural population of *Drosophila melanogaster*. Associative learning was tested in common-garden-raised flies collected in the spring and autumn over three consecutive years. Spring flies consistently learned better than autumn flies, revealing seasonal evolution of improved learning performance in nature. Fecundity showed the opposite seasonal pattern, suggesting a trade-off between learning and reproduction that also held at the level of individual females. This trade-off is mediated at least in part by natural polymorphism in the RNA-binding protein *couch potato* (*cpo*): a genotype favoured during summer showed poorer learning performance and higher fecundity than a genotype favoured over winter. Thus, seasonal environments can drive rapid cyclical evolution of learning performance, but the evolutionary dynamics may be driven by trade-offs generated by pleiotropic effects of causative alleles selected for other reasons.

## Introduction

Learning is a powerful mechanism of plasticity that allows animals to behaviourally respond to changing environments without genetically-based evolutionary change [1,2]. Learning is predicted to be favoured over fixed behavioural responses under a relatively narrow set of environmental conditions where cues are reliable, opportunities for learning are available, and the costs of errors are high [3]. Associative learning, the ability to link a stimulus with a positive or negative outcome, plays an important role in many aspects of animal biology including foraging, spatial orientation, predator avoidance, aggression, social interactions, and sexual behaviour [1,4–8] and has been shown to directly improve fitness under natural conditions [9]. More broadly, cognitive performance has been directly linked to fitness in several wild animal populations: memory ability predicts reproductive success in wild birds [10], spatial cognition is under natural selection in food-caching chickadees [11], and cognitive performance is associated with survival and longevity in both birds and primates [12,13], suggesting that variation in neural function has real fitness consequences in nature.

The ability to learn is itself a product of evolution, but the rate at which learning characteristics evolve in natural populations remains poorly understood. Examples of genetic differences in learning performance have been reported between closely related species [14,15] and between allopatric populations within a species [16–18], but the timescale over which these differences have evolved is unknown. Population differences in spatial memory and hippocampal structure that persist in a common garden environment have been documented along gradients of climate harshness in food-caching birds, demonstrating a genetic basis for cognitive variation in wild populations [19–21]. Laboratory selection experiments in a range of taxa indicate that natural populations have copious standing genetic variation for learning ability and that learning performance can improve rapidly under direct selection [22–27]. Natural variation in learning speed has also been documented within wild populations and linked to ecologically relevant outcomes. For example, faster-learning bumblebees show higher foraging efficiency in the field [4]. However, selection on learning in nature may be indirect and mediated by the fitness consequences of the many behaviours it modifies. An environmental change that results in an increased advantage of learning in the context of one behaviour (e.g., foraging) might have no effect on, or even decrease, its benefits in a different context (e.g., predator avoidance), thus dampening net selection on learning; alternatively, if multiple contexts shift in concert, selection could be amplified [3].

*D. melanogaster* is a well-established model for studying rapid evolution in a natural context, whose capacity for associative learning in laboratory settings is well characterised. As a multivoltine species producing 10–15 generations per year in temperate climates [28], successive generations experience a continuously shifting sequence of environmental conditions: populations pass through gradients of temperature, resource availability, population density, and pathogen pressure as the year progresses. These shifting conditions drive rapid, predictable, and repeatable cyclical evolution of life history, stress resistance, and immunity traits [28–31]. Population size and resource availability typically peak in summer as flies exploit abundant fruit resources and reproduce rapidly across several overlapping generations, whereas populations that persist through the colder months face resource scarcity, low density, and the physiological demands of overwintering, which favours traits such as reproductive dormancy and stress tolerance. Genomic analyses of seasonal allele frequency change across this annual cycle reveal that the resulting adaptation disproportionately involves genes associated with neural structure and function, suggesting that the neural architecture underlying behaviour is itself a primary target of this fluctuating selection [28]. One might therefore predict that learning ability, as an emergent property of neural circuit structure and function, should differ between spring and autumn populations. More specifically, the increased resource availability and social complexity of summer may intensify selection on learning through its role in foraging, pathogen avoidance, and sexual and social interactions [7,32–36], favouring higher learning ability in flies emerging after summer (i.e., autumn collections). Alternatively, learning may also contribute to overwintering-relevant behaviours such as shelter identification and cold-season foraging, and the evolutionary dynamics of learning ability may additionally be driven by selection on genetically correlated traits. Such genetic correlations may reflect the high energetic costs of neural tissue [37,38], generating trade-offs with other fitness traits. Experimental increases in brain size have been shown to reduce reproductive output in vertebrates [39,40], cognitive ability has been shown to influence life history variation in wild birds [41], and trade-offs between learning and reproduction have been reported in *Pieris rapae* [42] and *Drosophila* [43], as well as between learning and stress resistance in *Drosophila* [44,45]. Thus, seasonal evolution in learning ability could be driven by indirect effects of selection on correlated traits, in particular fecundity in the summer or stress tolerance in the winter.

Among the many genes that vary seasonally, *couch potato* (*cpo*) stands out as a particularly well-motivated candidate for mediating a seasonal learning-fecundity trade-off. *cpo* encodes a pleiotropic RNA-binding protein with documented effects on reproductive dormancy, fecundity, activity levels, stress resistance, and lifespan [46–48]. Multiple SNPs across the gene show strong and repeatable latitudinal clines and seasonal oscillations in allele frequency across North American and European populations [48–51], placing it among the most consistently identified targets of seasonal selection in this species. Critically, natural regulatory variation at *cpo* acts preferentially through neural tissue: head-tissue eQTL analyses identify 45 significant cis-eQTLs in head and none in body [48,52], and the protein is highly expressed in the nervous system including the mushroom body, the centre of associative learning in insects [53–55]. The concentration of regulatory variation in neural tissue, combined with *cpo*’s established role in seasonal adaptation and its pleiotropic effects on traits predicted to trade off with learning, generates a specific prediction: that the spring and autumn *cpo* genotypes should differ in learning performance in directions consistent with their seasonal frequency dynamics.

Here we test whether learning performance evolves rapidly over an annual timescale in a natural population of *D. melanogaster*. We assessed aversive and appetitive learning in common-garden-raised flies derived from spring and autumn collections from the same location over three consecutive years. We found that learning performance was consistently higher in the post-winter spring collections compared to the post-summer autumn collections. A reproductive cost of learning was indicated by an inverse relationship between learning performance and fecundity, observed both between seasonal populations and at the level of individual flies. Given that natural variants at *cpo* fluctuate in frequency with both season and latitude [28,46–48,56] and have documented effects on fecundity and reproductive dormancy [32–35], we reasoned that *cpo* is a plausible candidate for mediating this seasonal learning-fecundity relationship at the molecular level. We hypothesised that if *cpo* contributes to seasonal evolution in this way, flies carrying the spring-associated *cpo* genotype should mirror the seasonal differences and show higher learning and lower fecundity than those carrying the autumn-associated genotype. We found that learning performance and fecundity in flies homozygous for the spring (*cpo*^TTA^) versus autumn (*cpo*^CGT^) genotype closely paralleled the seasonal differences in the natural populations, suggesting that rapid fluctuating evolution of learning ability in wild *Drosophila* is at least partly driven by pleiotropic effects of *cpo* polymorphism on the neural substrate of learning.

## Methods

### (a) Seasonal populations derived from nature

The evolution of learning differences over seasonal time was assessed by comparing seasonal reconstructed populations derived from isofemale lines. Gravid females aspirated off decaying fruit in the spring (June) and autumn (November) at Linvilla Orchards in Media, PA (39.9°N, 75.4°W) across three consecutive years (2012–2014) were used to establish isofemale lines maintained in standard laboratory conditions (25°C, 12L:12D on cornmeal-molasses medium) on a four-week transfer cycle [28] (Figure 1a). Populations representing each collection were constructed after all collections were complete using 40 isofemale lines per collection (∼800 individuals per population at founding) and subsequently maintained in common garden culture for more than 10 non-overlapping generations. Systematic differences in traits between spring and autumn seasonal reconstructed populations assessed in the common garden are attributed to genetic differences in spring and autumn allele frequencies.

**Figure 1.**
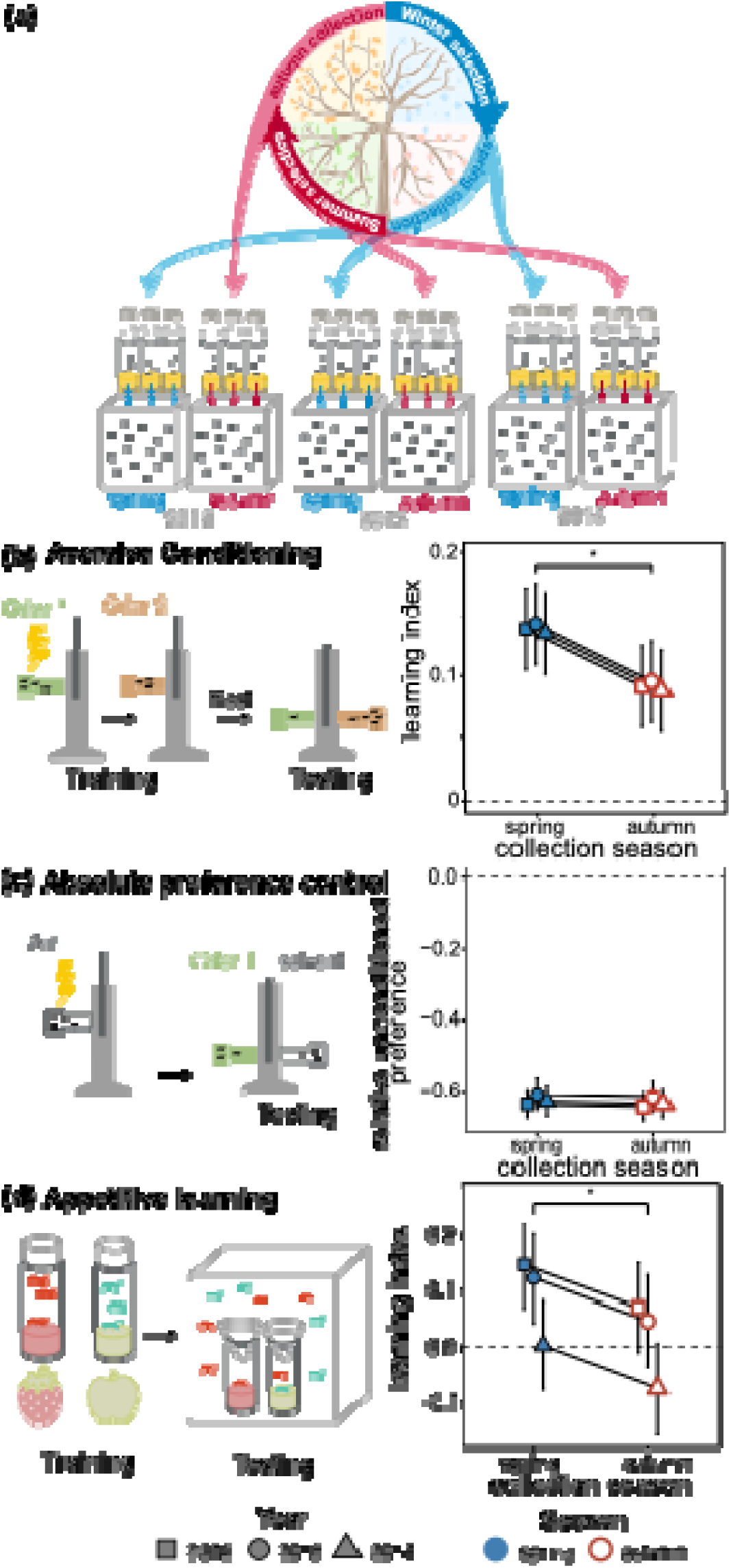
Rapid evolution of learning in natural populations of *Drosophila melanogaster*. (a) Seasonal collection scheme: gravid females were collected from spring and autumn populations at Linvilla Orchards over three consecutive years (2012–2014) and used to found common-garden reconstructed populations. (b) Aversive conditioning: spring populations showed higher learning (mean ± SE, model-based) than autumn populations, replicated across three years: 2012 (square), 2013 (circle), 2014 (triangle). Filled/blue symbols = spring; open/red symbols = autumn. Significance bracket indicates Season effect (p = 0.03). Values range from −1 (complete preference for the shock-paired odour) to 1 (complete preference for the safe, unpaired odour). (c) Absolute preference control: no difference in preference between odour and solvent in unconditioned flies. Values range from −1 (complete preference for solvent) to 1 (complete preference for the odorant). (d) Appetitive learning in outdoor field cages: spring populations returned to their conditioned flavour at a higher rate than autumn populations (p = 0.010). Values range from −1 (complete preference for the untrained flavour) to 1 (complete preference for the trained flavour).

### (b) cpo recombinant outbred populations

Three SNPs in *cpo* define the temperate (*cpo*^TTA^) and subtropical (*cpo*^CGT^) genotypes: two intronic SNPs (3R:17964408, 3R:17965558) and one SNP at 3R:17967866 (*D. melanogaster* reference genome v.6). The *cpo*^TTA^ genotype is more common at high latitudes and in the spring, while *cpo*^CGT^ is more common at low latitudes and in the autumn [48]. The SNP at 3R:17967866 was originally identified as a non-synonymous coding change in exon 5 with functional effects on reproductive dormancy and other life history traits [46–48,53,54]; however, its classification as a functional coding variant has been revised as it is not identified in the modENCODE catalogue of functional elements [57] and its molecular function remains unresolved [54]. Learning performance of the genotypes was assessed using ROPs each fixed for one genotype in a randomised genetic background [58]. ROPs for *cpo*^TTA^ and *cpo*^CGT^ were constructed using nine independent homozygous lines from the *Drosophila* Genetics Reference Panel [48] (Figure 3b). Ten gravid females from each line were permitted to lay eggs for 48 hours; after at least 10 non-overlapping generations of recombination among lines carrying the same genotype, each ROP was fixed for either *cpo*^TTA^ or *cpo*^CGT^, and any linked loci, in a heterogeneous outbred background[31,59]. All behavioural assays were conducted at 25°C under 12L:12D conditions.

**Figure 2.**
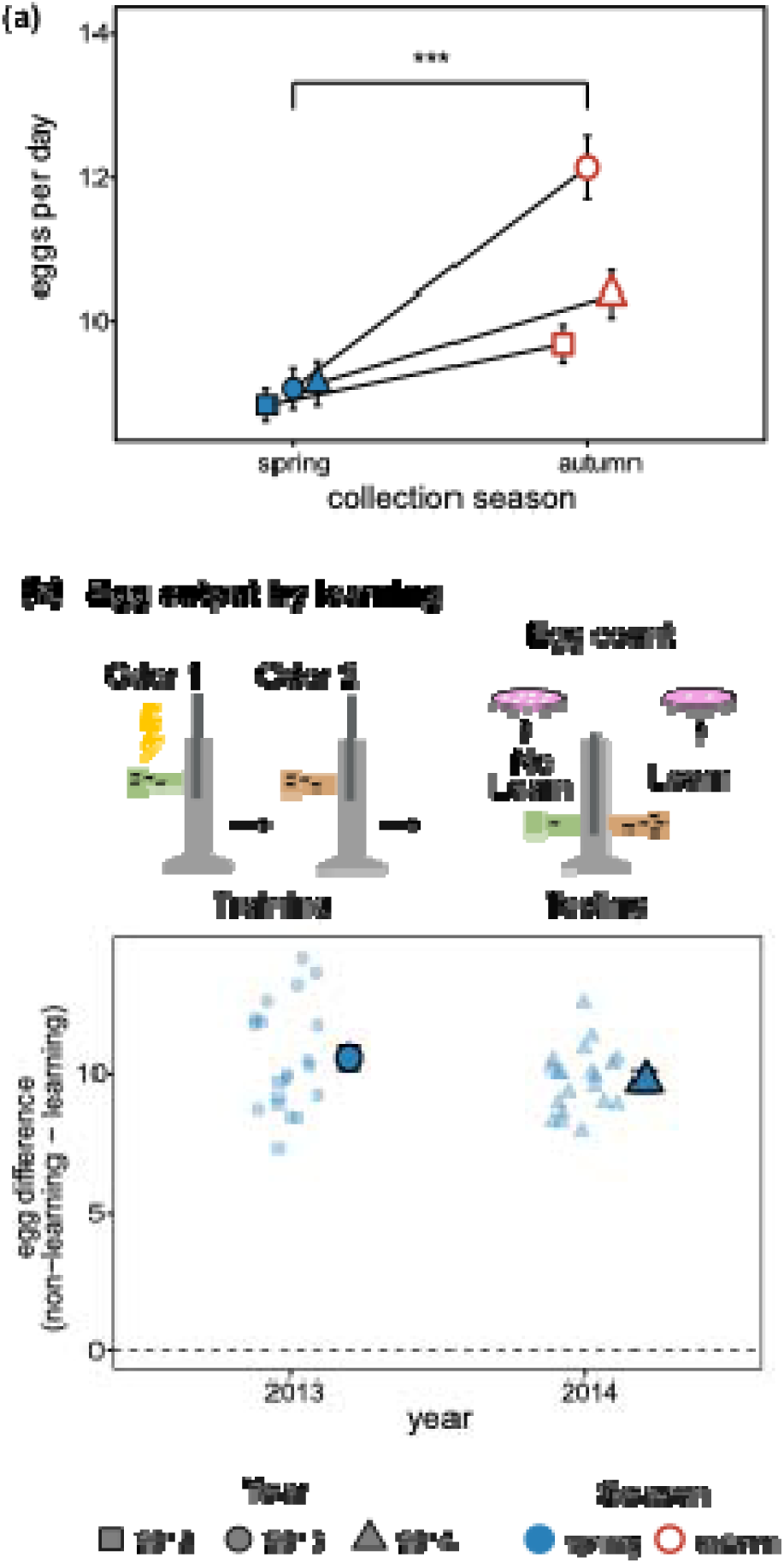
Fecundity trade-off in seasonal populations of *Drosophila melanogaster*. (a) Fewer eggs laid per day in the spring populations compared to the autumn, averaged over ten days; significance bracket indicates Season effect from linear model (p < 0.001). (b) Individuals that do not learn in the aversive conditioning assay (non-learners) lay more eggs per day over three days than those that learn, among replicates where the group showed net learning (paired t-test, t₁₄ = 2.84, p = 0.013). Small pale points show individual replicate differences; large filled points show mean difference ± SE by year.

**Figure 3.**
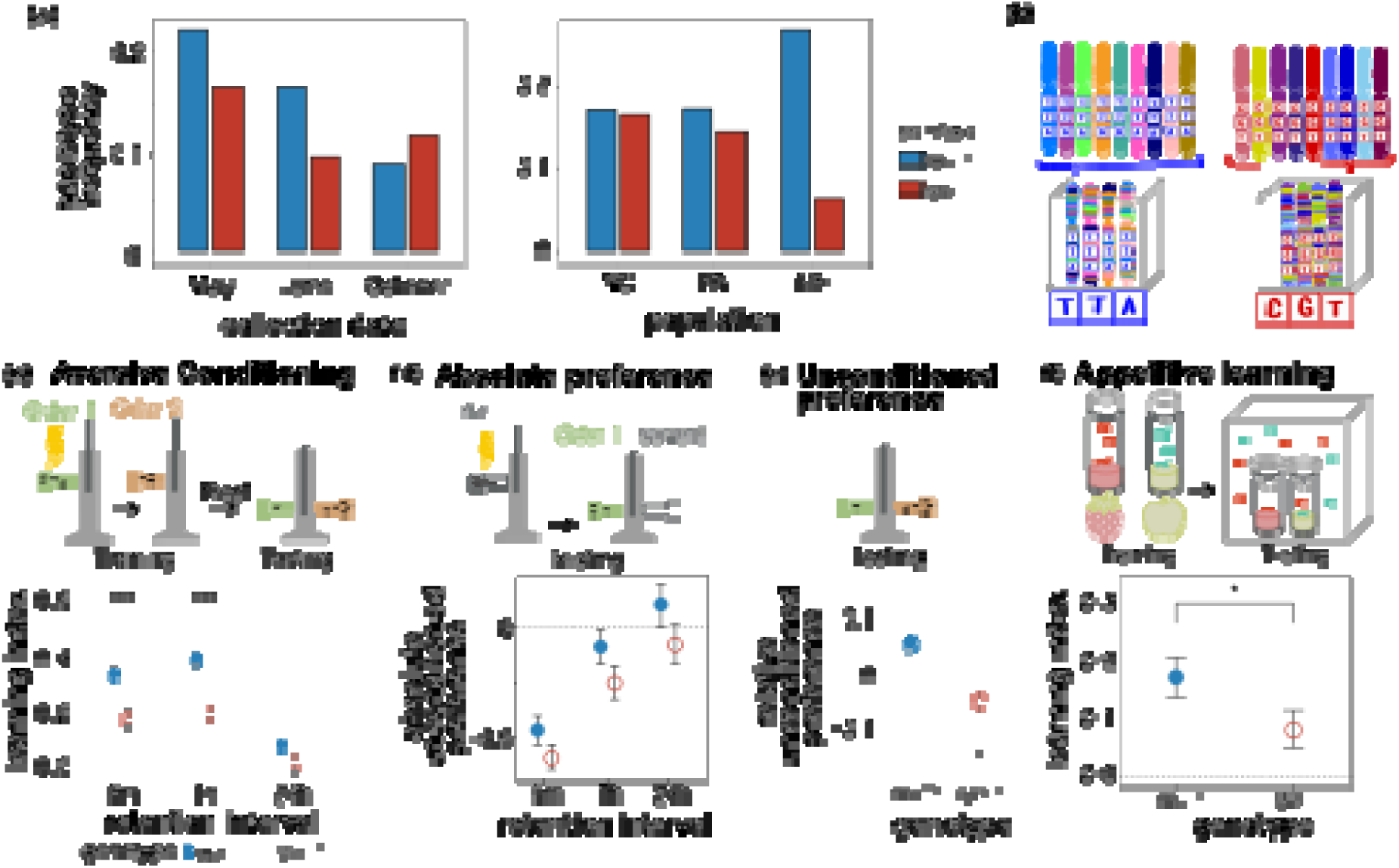
Natural variants in *couch potato* (*cpo*) show a learning difference paralleling the seasonal pattern. (a) Observed frequency of the *cpo*^TTA^ and *cpo*^CGT^ haplotypes across three collection timepoints (May, June, October) in a Pennsylvania orchard and across three populations along the latitudinal cline (right; North Carolina (NC, 35.5°N), Pennsylvania (PA, 39.9°N), Maine (ME, 44.0°N)) [48]. (b) The three SNPs defining the *cpo*^TTA^ and *cpo*^CGT^ haplotypes. Recombinant outbred populations (ROPs) were founded from independent inbred DGRP [92] strains each homozygous for one haplotype, then recombined over multiple generations to produce outbred populations fixed for the same three-SNP haplotype in a randomised genetic background [48]. (c) Aversive conditioning: flies homozygous for *cpo*^TTA^ showed higher learning (mean ± SE, model-based) than *cpo*^CGT^ across 5-minute and 1-hour retention intervals, with no difference at 24 hours. Values range from −1 (complete preference for the shock-paired odour) to 1 (complete preference for the safe, unpaired odour). (d) Absolute preference control: *cpo*^TTA^ flies showed slightly higher absolute odour avoidance than *cpo*^CGT^ at the 1-hour interval only. Values range from −1 (complete preference for solvent) to 1 (complete preference for the odorant). (e) Relative preference control: no difference in preference between the two experimental odours in unconditioned flies. Values range from −1 (complete preference for MCH) to 1 (complete preference for OCT). (f) Appetitive learning in outdoor field cages: *cpo*^TTA^ flies returned to their conditioned flavour at a higher rate than *cpo*^CGT^ flies. Values range from −1 (complete preference for the untrained flavour) to 1 (complete preference for the trained flavour).

### (c) Aversive shock learning in the laboratory

We quantified learning performance using an aversive olfactory conditioning assay in which flies learn to avoid an odour previously paired with mechanical shock [44]. Flies were reared in standard laboratory conditions (25°C, 12L:12D) at a standardised density of 100 eggs per vial, sorted into single-sex groups of 30 under light CO₂ anaesthesia 24 hours prior to assessment, and assayed at 3–5 days of age. Groups were conditioned to associate an odorant with mechanical shock across three conditioning cycles. Each cycle consisted of 30 seconds exposure to one odorant paired with mechanical shock (1-second pulses every 5 seconds), designated the conditioned stimulus predicting shock (CS+), followed by 60 seconds of humid air, 30 seconds exposure to a second odorant without shock, designated the unpaired conditioned stimulus (CS−), and 60 seconds of humid air. Methylcyclohexanol (MCH, 800 µL/L) and 3-octanol (OCT, 600 µL/L) served as odorants; conditioning direction was counterbalanced across groups so that each odorant served as CS+ in half the groups, controlling for innate odour preference. Memory was tested after a one-hour retention interval by allowing flies 60 seconds to choose between the two odorants in an elevator T-maze. The *cpo* genotypes were additionally assessed at 5-minute (immediate memory) and 24-hour (long-term memory) retention intervals.

Two types of control experiments excluded differences in odour detection or innate preference as explanations for learning differences. Controls for absolute preference subjected flies to mechanical shock without odorant exposure before offering a choice between a single odorant and clean air; this control was conducted for both the seasonal populations and the *cpo* genotypes. Controls for relative preference assessed innate odour preference in naïve flies choosing between the two odorants; this control was conducted only for the *cpo* genotypes. Both controls were assessed using the same GLMM framework described above, with Replicate block as a random effect. Absolute preference modelled the proportion of flies choosing the odorant over solvent, as a function of Year (seasonal populations) or Genotype (*cpo*), Sex, and Odorant identity. Relative preference modelled the proportion of flies choosing OCT over MCH, as a function of Genotype, Sex, and Odorant identity.

All statistical analyses were performed in R (v4.3; R Core Team 2024), with Claude (Anthropic) assisting with debugging and reformatting analysis code; all statistical analyses, results, and interpretations were performed and verified by the authors. Learning performance was modelled as the proportion of flies selecting unconditioned stimulus (CS-) using generalised linear mixed models (GLMMs) with a logit link and binomial error distribution, fitted by maximum likelihood (lme4 v1.1; Bates et al. 2015). Significance of fixed effects was assessed using likelihood ratio tests (afex v1.3; [60]). Flies remaining in the centre of the T-maze were excluded. For visualisation, the proportion of flies choosing the unpaired odorant (CS-) was rescaled as 2 x proportion − 1, yielding a learning index ranging from −1 (complete preference for the shock-paired odorant, CS+, indicating no avoidance) to 1 (complete preference for the unpaired odorant, CS-, indicating maximal avoidance learning), where 0 indicates no preference and thus no learning. Model-based standard errors from estimated marginal means (emmeans v1.8; [61]) are shown in all figures. Replicate block was included as a random effect in all models. Fixed effects structures were:

*Natural populations:* Proportion ∼ Year + Season + Sex + Odorant + Year:Odorant+ Sex:Odorant + (1 | Block)

*cpo genotypes:* Proportion ∼ Genotype + Sex + Odorant + Retention interval + Genotype:Odorant + Genotype:Retention interval + (1 | Block)

### (d) Appetitive learning in a natural environment

To test whether learning differences among genotypes observed in the controlled laboratory environment translate to ecologically relevant behaviour in a complex, semi-natural setting, we assessed appetitive learning as the proportion of flies caught in traps flavoured with the same flavour as their conditioning food. The assay was adapted from [25,35] and conducted in outdoor field cages over two replicate days (30–31 August 2016) at the University of Pennsylvania.

Flies were reared under standard conditions (25°C, 12L:12D) at controlled density (∼200 eggs per bottle) and sorted into single-sex groups of 50 at 3 days of age. Groups were marked with fluorescent powder in one of four colours (Blue 1162B, Yellow 1162Y, White 1162W, Pink 1162P; Bioquip, Rancho Dominguez, CA) according to source population and conditioning direction; a pinch of powder was added to an empty vial, flies were added and gently rolled until coated. Flies were transferred to non-nutritional agarose for 12 hours to recover and become food-motivated, then exposed to either strawberry- or apple-flavoured food for an 8-hour conditioning period, followed by a 4-hour retention period on plain agar, before release into outdoor cages. The four-colour marking scheme permitted simultaneous release of flies from two populations (spring and autumn, or *cpo*^TTA^ and *cpo*^CGT^) conditioned to each flavour (∼400 flies per conditioning direction per cage). Mesh cages (0.6 × 0.6 × 1.8 m) with plant-bedded floors were positioned beneath a peach tree, each containing four pairs of unidirectional strawberry and apple traps. Flies caught within 14 hours of release were sexed and assigned to population and conditioning direction by colour. All recovered flies retained powder markings, confirming effective marking and no infiltration from outside. Statistical models followed the same structure as for aversive conditioning, with trap and day as nested random effects:

*Natural populations:* Proportion ∼ Year + Season + Sex + CS + Year:CS + Sex:CS + (1 | Day:Trap)

*cpo genotypes:* Proportion ∼ Genotype + Sex + CS + Genotype:Sex + (1 | Day:Trap)

### (e) Fecundity

Population-level fecundity was measured by placing 25 virgin females from each ROP with 25 males from a standardised tester stock (BDSC-3605) in culture bottles. Food plates were replaced daily for ten days to count eggs laid and record mortalities; daily mean fecundity per surviving female was calculated across 10 replicate bottles per population. For seasonal populations, daily fecundity was averaged within each bottle across the ten-day assay period, and bottle means were compared between populations using a linear model with season and year as fixed effects. For *cpo* ROPs, genotype was the fixed effect. Significance was assessed using F-tests.

The within-population relationship between learning and fecundity was assessed in spring ROPs from 2013 and 2014. Virgin females were aged for 3 days, then subjected to the aversive olfactory conditioning and test described above, and classified as learners (chose CS−) or non-learners (chose CS+). From each replicate test where the group’s aggregate T-maze choice showed net learning (learn_freq ≥ 0), five learner and five non-learner females were each housed with 5 males from a tester stock on yeast-supplemented food to stimulate egg production, then transferred to grape-agar for egg collection. Daily fecundity was recorded for 3 days and mean eggs per female per day was calculated accounting for any female mortality. Because learners and non-learners were drawn from the same conditioning replicate under identical conditions, a paired t-test was used to account for between-replicate variation in absolute fecundity, treating each replicate test as a matched pair.

## Results

### (a) Rapid evolution of learning and fecundity in a natural population

Learning ability in the wild *D. melanogaster* population evolved rapidly and repeatedly across seasonal time. Spring collections showed higher learning than autumn collections from the same year when assessed using laboratory aversive conditioning (χ^2^_1_ = 4.68, p = 0.03; Figure 1b, Supplemental Table 1). These behavioural differences were not attributable to differences in odour perception, as there was no effect of season on absolute preference between odorant and solvent in unconditioned flies ( ^2^ = 0.03, p = 0.85; Figure 1c).

This seasonal difference in learning was not limited to the laboratory aversive assay. In outdoor mesocosms, spring flies were also more likely than autumn flies to be caught in traps flavoured with the same flavour as the food they had previously encountered during appetitive conditioning (χ^2^_1_ = 6.66, p = 0.010; Figure 1d), demonstrating that the seasonal learning difference generalises across conditioning paradigms and testing contexts.

The seasonal pattern of fecundity was opposite to that of learning ability. At the population level, autumn populations laid on average 30% more eggs per female per day than spring populations (F_1,55_ = 35.72, p = 1.76×10^-7^; Figure 2a). A phenotypic trade-off between learning and reproduction was also apparent at the individual level within the spring populations. Females that chose the shock-associated odour (CS+, i.e., non-learners) subsequently laid an average of eight more eggs per day (approximately 23% more) than females that avoided the shock-associated odour (CS−, i.e., learners) from the same replicate test (t_14_ = −2.84, p = 0.013; Figure 2b). The inverse relationship between learning performance and fecundity thus holds both among populations differing in seasonal origin and among individuals within a population.

### (b) Natural variation in cpo sequence affects learning and fecundity

The latitudinal and seasonal frequency patterns of the *cpo*^TTA^ and *cpo*^CGT^ genotypes show opposite trends: *cpo*^TTA^ increases in frequency with latitude and is more common in spring collections, while *cpo*^CGT^ shows the opposite pattern [48] (Figure 3a). Recombinant outbred populations (ROPs) homozygous for the spring-associated *cpo*^TTA^ genotype showed higher aversive learning than ROPs fixed for the autumn-associated *cpo*^CGT^ genotype, assessed across 5-minute, 1-hour, and 24-hour retention intervals (χ^2^_1_ = 27.12, p = 1.91 × 10^-7^; Figure 3c, Supplemental Table 1). At the 24-hour retention interval, *cpo*^CGT^ flies showed no memory above chance (p = 0.602), while *cpo*^TTA^ flies retained a marginal memory trace (p = 0.059), suggesting that the spring genotype confers greater persistence of memory consolidation. The *cpo* ROPs did not differ in the relative odour preference of naïve flies choosing between the two odorants (χ^2^_1_ = 1.11, p = 0.29; Figure 3e), although there was a genotype-by-odorant interaction (χ^2^ = 6.24, p = 0.012), with the *cpo*^TTA^ ROP showing a slight preference for MCH and the *cpo*^CGT^ ROP for OCT. Because flies were conditioned in both directions to avoid either odour, this slight asymmetry in innate preference does not affect the quantification of learning ability. In the absence of conditioning, *cpo*^TTA^ flies showed stronger avoidance of odorants than *cpo*^CGT^ flies at the 1-hour interval (z = 2.88, p = 0.004; Figure 3d) but not at 5-minute (z=1.37, p=0.17) or 24-hour (z=0.48, p=0.63) intervals. This difference in absolute odour avoidance likely reflects a genotype difference in odour sensitivity or general arousal state rather than a learned association and is unlikely to inflate learning scores given the counterbalanced conditioning design.

The spring-associated *cpo*^TTA^ ROP also showed higher appetitive associative learning in the natural mesocosm assay (χ^2^_1_ = 3.72, p = 0.054; Figure 3f). This result is broadly consistent with the genotype effect on learning generalising beyond the laboratory aversive paradigm, though this result did not reach conventional significance. Consistent with the population-level trade-off observed in the seasonal ROPs, *cpo*^TTA^ females laid 17% fewer eggs per day than *cpo*^CGT^ females, though this difference was marginal given the limited number of replicate lines (F_1,18_ = 3.99, p = 0.061; Figure 4). Together, these results indicate that the *cpo* haplotype recapitulates the seasonal phenotypic differences observed in natural populations, linking natural variation at this genomic region to the rapid cyclical evolution of both learning ability and fecundity in wild *Drosophila*, and implicating *cpo* itself, or variants in strong linkage disequilibrium, as pleiotropic mediators of this seasonal phenotypic cycle.

**Figure 4.**
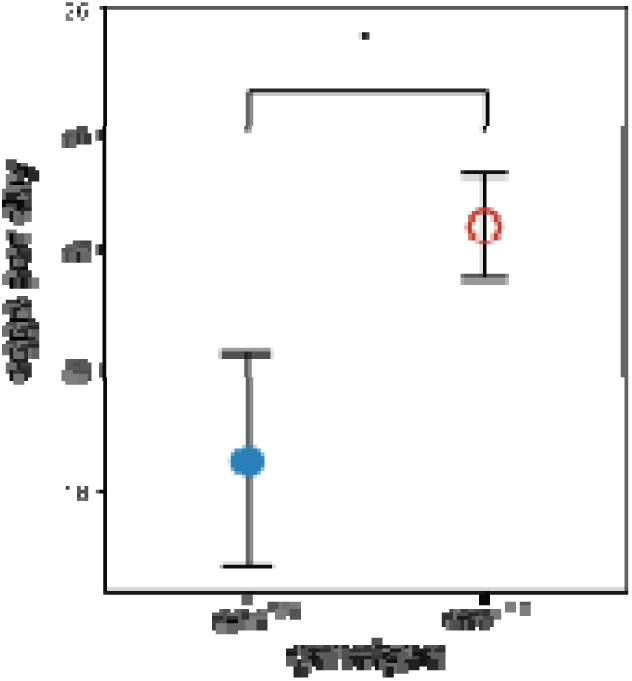
Fecundity by *cpo* genotype. Flies homozygous for the autumn-associated *cpo*^CGT^ genotype laid more eggs per day than flies homozygous for the spring-associated *cpo*^TTA^ genotype.

## Discussion

### (a) Rapid evolution in natural populations

Our data show that learning and fecundity both evolve rapidly in a natural population of *D. melanogaster* over approximately 10–15 generations from spring to autumn and 1–2 generations from autumn to spring [28]. Because flies were reared and tested in common laboratory conditions at 25°C, environmental effects are excluded and the observed differences can be attributed to genetic changes cycling on an annual basis. The repeatability of these differences across years rules out genetic drift as an explanation, implicating instead deterministic evolutionary processes. Genomic evidence strongly suggests that gene flow through migration does not generate the observed seasonal cycles in allele frequency [51,62]; moreover, genome-wide analyses of seasonal adaptation reveal that genes associated with neural structure and function, including axon guidance, neuron projection morphogenesis, and CNS development, are among the most strongly and repeatably enriched targets of seasonal selection across years and populations [28]. The rapid and repeatable seasonal divergence in learning performance we report here thus has a clear genomic counterpart, and demonstrates that learning ability is not only a mechanism of phenotypic plasticity allowing organisms to respond to environmental change, it is itself a likely target of rapid adaptation in wild populations [11,12,63].

The learning differences between spring and autumn populations were consistently observed in both a laboratory aversive conditioning assay and an appetitive conditioning assay conducted in outdoor mesocosms in late summer, with spring-origin flies outperforming autumn-origin flies in both contexts. Appetitive and aversive associations are formed at least partly in different subsets of mushroom body neurons and involve different neurotransmitters, yet both forms of memory depend on shared molecular mechanisms of synaptic plasticity in the mushroom body [64]. The parallel seasonal pattern across both assays therefore implies that the evolutionary change affected a shared component of the learning process upstream of the specific memory circuits, consistent with an experimental evolution study in which populations selected for appetitive learning also showed improved aversive shock-odour performance [65].

The higher learning in spring populations is counter to the intuitive prediction that learning should be most advantageous during summer, when flies are behaviourally active and engage in foraging, social, and sexual interactions known to depend on learning. However, cognitive abilities may evolve to match the demands of an organism’s biological and physical environment more broadly [66,67], and energetically challenging environments such as harsh winters are hypothesised to favour cognitive investment at the cost of other physiological systems [68]. Consistent with this, genome-wide analyses of seasonal adaptation in the same population reveal that neural structure and function genes are the most consistently and repeatably enriched targets of winter selection across years and environmental axes — more so than stress tolerance or metabolic genes — suggesting that selection on neural architecture is a general feature of winter adaptation rather than an incidental correlate [28]. The spring learning advantage we document may thus reflect direct selection on neural function during winter rather than an indirect correlated response to selection on stress resistance or fecundity alone.

This interpretation is supported by compelling evidence from food-caching birds: population differences in spatial memory and hippocampal volume persist in common garden conditions along gradients of environmental harshness, with individuals from harsher climates showing superior learning performance [19,20], and hippocampal volume and neuron number increase along gradients of climate severity [69]. Furthermore, spatial cognitive ability has been directly linked to survival and longevity in wild chickadee populations [12], demonstrating that these cognitive differences have real fitness consequences in harsh environments. Learning may be important for overwintering-relevant behaviours in harsh climates such as locating suitable overwintering sites or navigating temperature-dependent foraging landscapes [70], analogous to the role of spatial learning in cache recovery in overwintering birds [71–73]. Alternatively, seasonal evolution of learning may be driven not by direct selection on learning itself, but by selection on genetically correlated traits such as fecundity and stress resistance, a possibility we consider directly below.

### (b) Links between resilience, fecundity and learning

Autumn collections showed a genetic component to seasonal changes in reproductive output with higher fecundity than spring collections. This seasonal fecundity advantage in autumn populations is consistent with strong selection for rapid reproduction during summer months of population expansion [74]. These fecundity differences sit within a broader pattern of seasonal adaptation. Spring-derived lines show greater propensity for reproductive dormancy [29], higher resistance to thermal stress and starvation [28,30], and stronger resistance to bacterial pathogens [31], a suite of resilience traits expected to be favoured by harsh winter conditions and known to trade off with fecundity [75,76]. The present results reinforce the hypothesis of alternating seasonal selection: for resilience traits during winter and for reproductive output during summer expansion [59,60].

Crucially, the inverse relationship between learning and fecundity was not limited to the population level. Within the spring populations, females that successfully avoided the shock-associated odour (learners) laid significantly fewer eggs over the subsequent three days than non-learners from the same replicate test. This individual-level trade-off is particularly informative because it cannot be explained by population-level genetic or environmental confounds: both learners and non-learners were drawn from the same replicate group under identical conditions, and the paired design explicitly accounts for between-replicate variation in baseline fecundity. The finding adds to a growing body of evidence that the costs of learning and memory generate genuine fitness trade-offs in natural contexts [42]. Cognitive ability has been shown to influence reproductive life history variation directly in wild birds [41], experimentally increasing brain size in guppies reduces reproductive output [40], and natural variation in learning speed in bumblebees is associated with differences in foraging returns [4]. A fitness cost of the learning ability itself distinct from the cost of individual memory traces has been demonstrated directly in *Drosophila* [45], and artificial selection for learning ability reduces competitive fitness [25]. Taken together, these results are consistent with previously reported trade-offs between learning and reproduction across taxa [42,43,77] and support the hypothesis that the seasonal decline in learning from spring to autumn reflects, at least in part, selection for increased fecundity during summer. If so, the concerted annual evolutionary changes in learning performance, fecundity, and resilience traits are jointly mediated by seasonal fluctuations in allele frequencies at loci with pleiotropic effects across all these traits.

It is important to note that both learning and fecundity were assessed here under optimal laboratory conditions with stable temperature, abundant high-calorie food, and no requirement for cognitive performance to obtain resources. The bountiful lab environment may not reflect the fitness dynamics operating in natural field environments. Under natural conditions, the ability to learn may itself contribute to foraging success, predator avoidance, and mate acquisition, such that the fitness cost of higher learning ability could be substantially smaller, or the benefit substantially larger, than our common-garden assays suggest. The trade-off we observe may therefore reflect the outcome of selection under natural seasonal conditions rather than an intrinsic physiological constraint, and the relative costs and benefits of learning are likely to shift across the seasonal environmental gradient in ways that our laboratory measures cannot fully capture.

### (c) Candidate polymorphism at cpo

Our results identify *cpo* as a strong candidate for one such pleiotropic locus mediating the annual seasonal evolutionary cycle. The *cpo*^TTA^ genotype increases in frequency with latitude and is more common in spring collections, while *cpo*^CGT^ shows the opposite pattern, consistent with the frequency trajectories expected under seasonally fluctuating selection [28,46,47,49–51,62]. Natural *cpo* variants have documented effects on activity, dormancy, fecundity, and lifespan [46–48], and the ROP design with randomised genetic backgrounds used here minimises the contribution of linked loci, strengthening the inference that the *cpo* locus itself, or variants in strong linkage disequilibrium, drives the observed phenotypic differences.

The *cpo*^TTA^ ROP showed higher learning across all retention intervals and in the natural mesocosm assay, closely mirroring the seasonal pattern in the natural populations. Fecundity was also lower in the *cpo*^TTA^ ROP than in the *cpo*^CGT^ ROP, with females laying approximately 17% fewer eggs per day (F_1,18_= 3.99, p = 0.061). The direction and magnitude of the effect are nonetheless consistent with the seasonal fecundity difference and with the broader trade-off pattern, and the convergence of learning, fecundity, and natural frequency data across multiple lines of evidence strengthens the case for *cpo* as a pleiotropic mediator of the seasonal phenotypic cycle.

That *cpo* affects learning is a new finding, though not entirely unexpected: the gene is expressed in the mushroom bodies and *cpo* loss-of-function mutants exhibit behavioural phenotypes [53,54]. Notably, *cpo* knockdown reduces odour avoidance [53], consistent with the stronger absolute odour avoidance we observed in the spring *cpo*^TTA^ ROP at the 1-hour interval. However, impaired odour detection would be expected to reduce rather than enhance learning performance, and the *cpo* ROPs did not differ in relative innate odour preference, arguing against a perceptual explanation for the learning difference. It is more plausible that the natural *cpo* variants studied here influence the neural processes of memory formation, storage, or recall. RNA-binding proteins are known to regulate local translation at synapses and play important roles in synaptic plasticity and memory consolidation [78–80], suggesting a plausible mechanistic route through which *cpo* variation could affect memory without altering sensory perception. The specific mechanism through which natural *cpo* variants modulate neuronal function remains to be determined.

### (d) Pleiotropy as a driving force in the evolution of learning

Discussions of learning evolution have typically emphasised ecological factors that modulate the net benefits of learning: environmental predictability, cue reliability, opportunity costs, and the consequences of errors [1,67,81,82]. Theory predicts that learning is most likely to be favoured when environments are variable but not unpredictably so, when reliable cues are available, and when the costs of learning are low relative to its benefits [3]. Our findings highlight a complementary and underappreciated force: natural genetic variants affecting learning may be embedded in highly pleiotropic loci whose overall fitness effects span behaviour, reproduction, and stress physiology simultaneously [83]. This is not unique to *cpo*; the broad pleiotropic effects of learning genes are well documented from mutant studies [83–86] and natural polymorphisms. A parallel case is the *foraging* (*for*) polymorphism in *D. melanogaster*, which affects not only learning [65,87] but also foraging strategy [88], aggregation [89], and competitive ability [87]. The genetic architecture of olfactory behaviour in *Drosophila* more generally involves hundreds of loci with broad pleiotropic effects on multiple behavioural and physiological traits [86], suggesting that the entanglement of learning with other fitness components may be a general feature of its genetic basis rather than a special property of *cpo*. Recent genomic work in food-caching chickadees has identified specific genes and gene networks underlying spatial cognitive variation in wild populations [21,90], opening the possibility of directly connecting genomic variation to cognitive phenotypes and fitness in natural systems, an approach that could powerfully complement the population-level evolutionary framework we employ here.

The broad phenotypic reach of such loci means that the net selection acting on them in natural populations reflects their aggregate impact across all affected traits, making it difficult to disentangle which phenotypes are the primary targets of selection and which evolve as correlated responses. In our system, we cannot determine from the present data alone whether seasonal changes in learning performance are driven by ecological selection on learning per se, or whether learning evolves primarily as a pleiotropic consequence of selection acting on fecundity, stress resistance, or other correlated traits. These need not be mutually exclusive; the same allele frequency change at *cpo* could simultaneously be favoured for its effects on dormancy in winter and generate a correlated improvement in learning as a mechanistic consequence of its action on neural tissue. Intriguingly, genome-wide analyses of seasonal adaptation suggest that genes associated with neural structure and function may be disproportionately represented among the targets of seasonal selection [28], which would be consistent with at least some of the learning evolution we observe reflecting direct selection on neural architecture rather than an indirect correlated response alone, although this remains to be tested directly. Resolving the relative contributions of direct and indirect selection will require connecting individual fitness consequences of learning differences to specific seasonal environments, a challenge that emerging tools linking genomic variation to cognitive phenotypes in wild populations [21] may help address. Regardless of the proximate cause, the learning differences between spring and autumn populations, and between the *cpo* genotypes, are likely to have ecologically meaningful consequences for behaviour in the wild, and illustrate how the rapid, predictable genomic turnover characteristic of seasonal adaptation can propagate through the nervous system to shape cognitive phenotypes on a rapid timescale.

## Acknowledgements

This work was supported by NSF GRFP DGE-0822 (ELB), the Rosemary Grant Award from the Society for the Study of Evolution (ELB), the Teece Dissertation Research Fellowship from the University of Pennsylvania (ELB), the Peachey Environmental Fund from the Department of Biology, University of Pennsylvania (ELB), and NIH R01GM100366 (PS), as well as research funding from the University of Lausanne to TJK. This work would not have been possible without the community supportive resources available at FlyBase [91].

## Author contributions

ELB, TJK and PS designed the experiment. ELB performed the experiment and conducted the analyses. ELB wrote the manuscript with feedback from TJK and PS.

---

**Supplemental Table 1.**
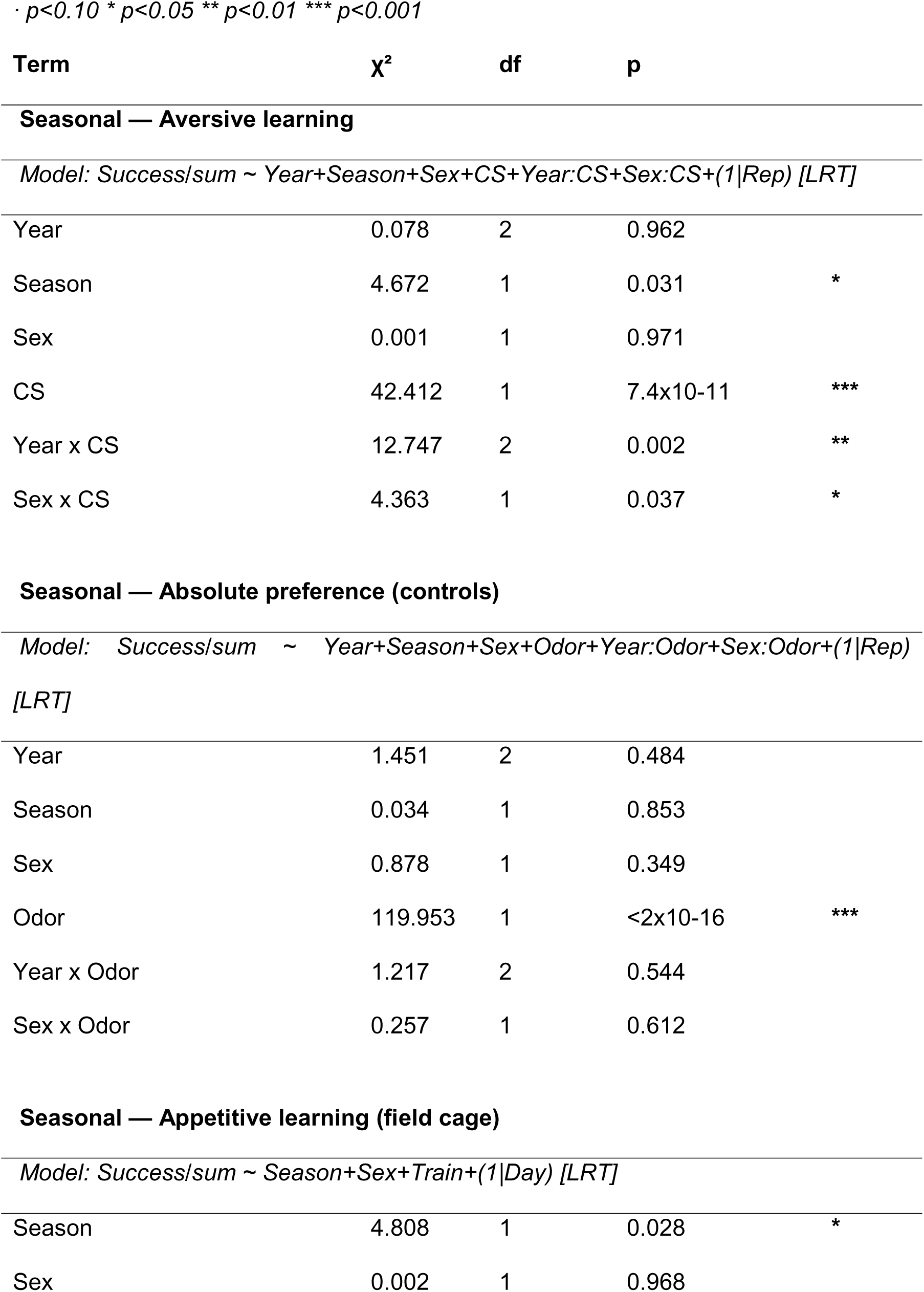
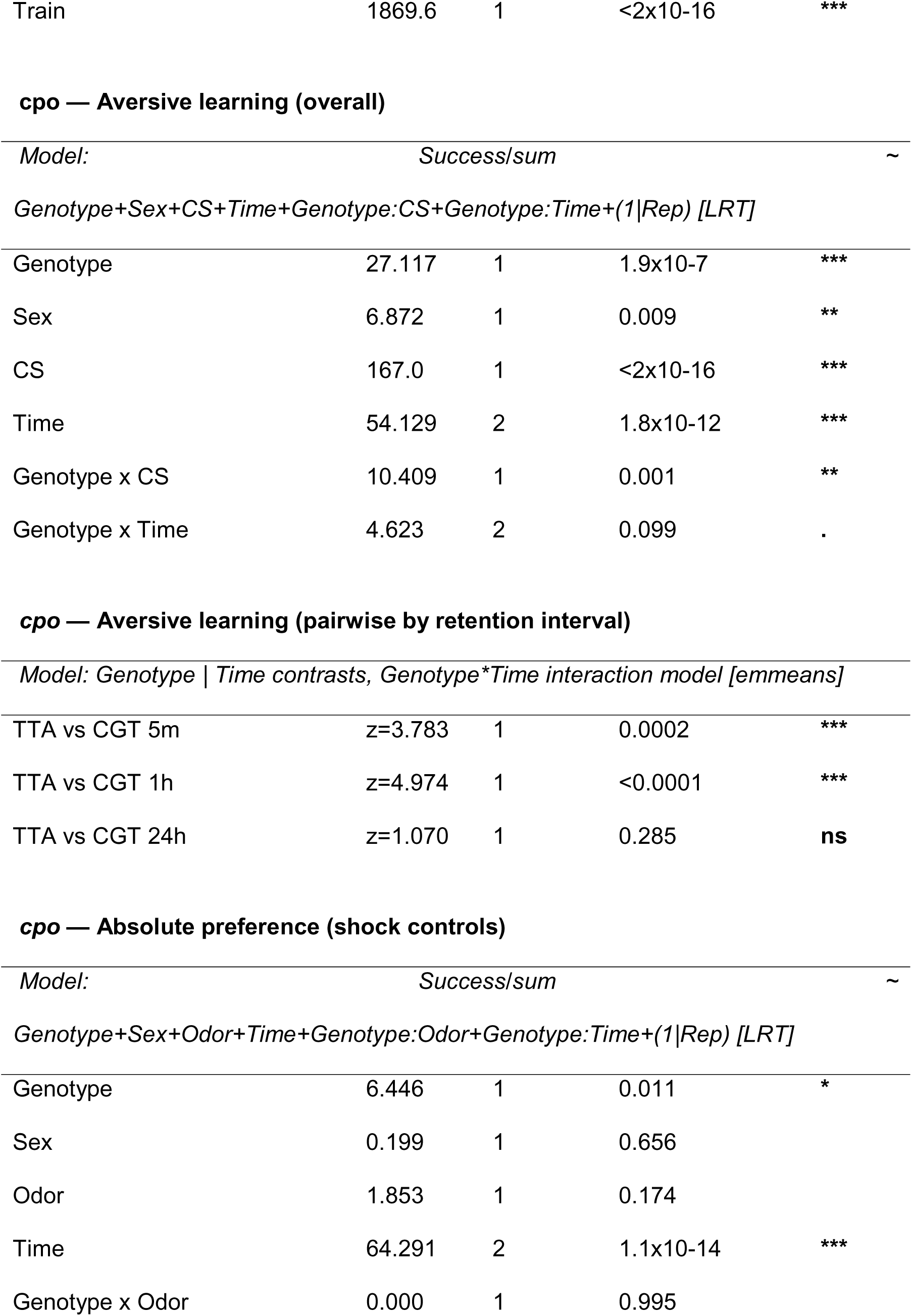
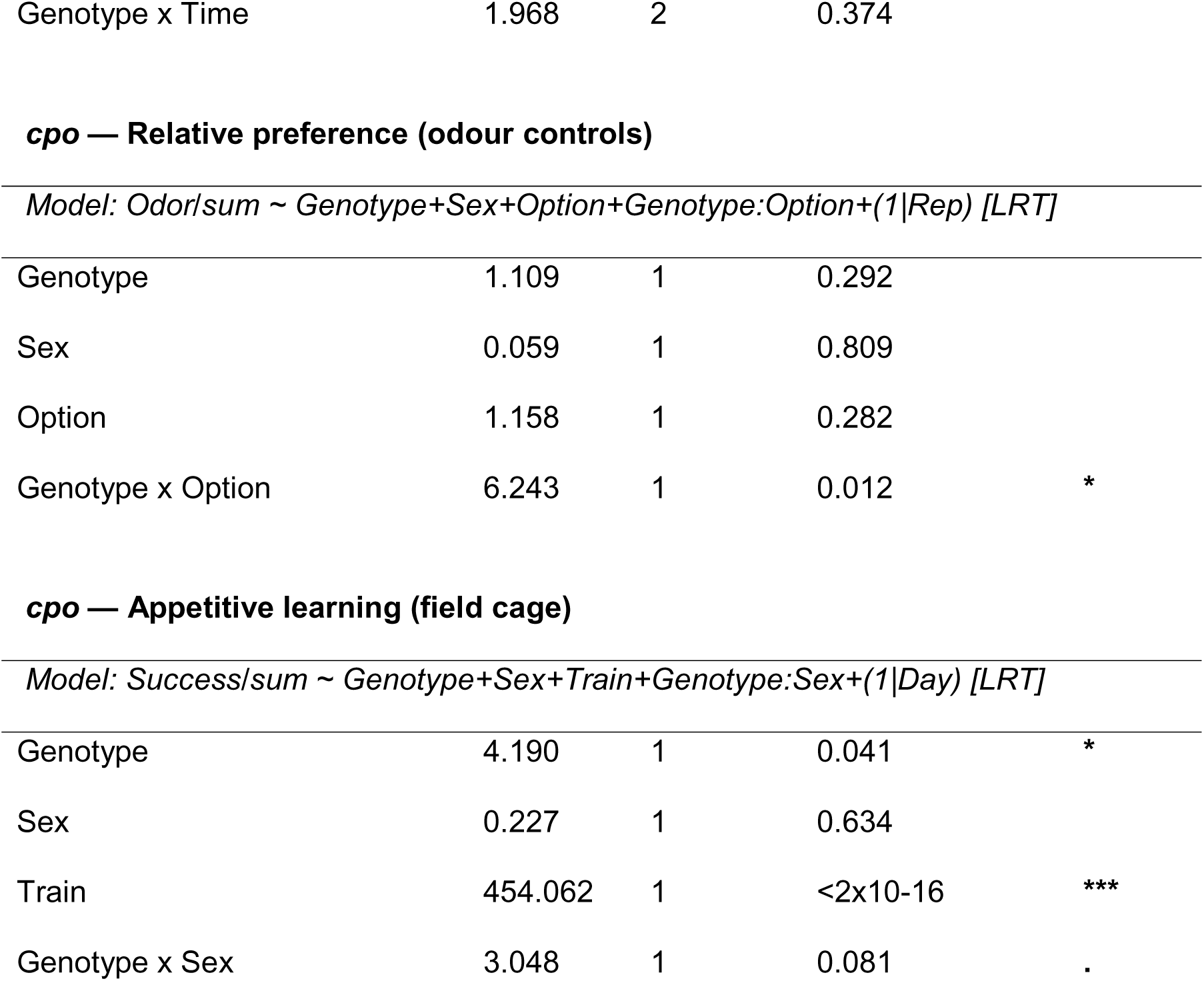
Mixed Model ANOVA.

· $p < 0.10$ \* $p < 0.05$ \*\* $p < 0.01$ \*\*\* $p < 0.001$
| Term | $\chi^2$ | df | p | |
| --- | --- | --- | --- | --- |
| **Seasonal — Aversive learning** |  |  |  |  |
| *Model: Success/sum ~ Year+Season+Sex+CS+Year:CS+Sex:CS+(1\|Rep) [LRT]* |  |  |  |  |
| Year | 0.078 | 2 | 0.962 |  |
| Season | 4.672 | 1 | 0.031 | * |
| Sex | 0.001 | 1 | 0.971 |  |
| CS | 42.412 | 1 | 7.4x10-11 | *** |
| Year x CS | 12.747 | 2 | 0.002 | ** |
| Sex x CS | 4.363 | 1 | 0.037 | * |
| **Seasonal — Absolute preference (controls)** |  |  |  |  |
| *Model: Success/sum ~ Year+Season+Sex+Odor+Year:Odor+Sex:Odor+(1\|Rep) [LRT]* |  |  |  |  |
| Year | 1.451 | 2 | 0.484 |  |
| Season | 0.034 | 1 | 0.853 |  |
| Sex | 0.878 | 1 | 0.349 |  |
| Odor | 119.953 | 1 | <2x10-16 | *** |
| Year x Odor | 1.217 | 2 | 0.544 |  |
| Sex x Odor | 0.257 | 1 | 0.612 |  |
| **Seasonal — Appetitive learning (field cage)** |  |  |  |  |
| *Model: Success/sum ~ Season+Sex+Train+(1\|Day) [LRT]* |  |  |  |  |
| Season | 4.808 | 1 | 0.028 | * |
| Sex | 0.002 | 1 | 0.968 |  |
| Train | 1869.6 | 1 | <2x10-16 | *** |

**cpo — Aversive learning (overall)**
| *Model:* | *Success/sum* |  |  | *~* |
| --- | --- | --- | --- | --- |
| *Genotype+Sex+CS+Time+Genotype:CS+Genotype:Time+(1\|Rep) [LRT]* |  |  |  |  |
| Genotype | 27.117 | 1 | 1.9x10-7 | *** |
| Sex | 6.872 | 1 | 0.009 | ** |
| CS | 167.0 | 1 | <2x10-16 | *** |
| Time | 54.129 | 2 | 1.8x10-12 | *** |
| Genotype x CS | 10.409 | 1 | 0.001 | ** |
| Genotype x Time | 4.623 | 2 | 0.099 | . |

**cpo — Aversive learning (pairwise by retention interval)**
| *Model: Genotype \| Time contrasts, Genotype\*Time interaction model [emmeans]* |  |  |  |  |
| --- | --- | --- | --- | --- |
| TTA vs CGT 5m | z=3.783 | 1 | 0.0002 | *** |
| TTA vs CGT 1h | z=4.974 | 1 | <0.0001 | *** |
| TTA vs CGT 24h | z=1.070 | 1 | 0.285 | ns |

**cpo — Absolute preference (shock controls)**
| *Model:* | *Success/sum* |  |  | *~* |
| --- | --- | --- | --- | --- |
| *Genotype+Sex+Odor+Time+Genotype:Odor+Genotype:Time+(1\|Rep) [LRT]* |  |  |  |  |
| Genotype | 6.446 | 1 | 0.011 | * |
| Sex | 0.199 | 1 | 0.656 |  |
| Odor | 1.853 | 1 | 0.174 |  |
| Time | 64.291 | 2 | 1.1x10-14 | *** |
| Genotype x Odor | 0.000 | 1 | 0.995 |  |

|  |  |  |  |
| --- | --- | --- | --- |
| Genotype x Time | 1.968 | 2 | 0.374 |
| Genotype | 1.109 | 1 | 0.292 |  |
| Sex | 0.059 | 1 | 0.809 |  |
| Option | 1.158 | 1 | 0.282 |  |
| Genotype x Option | 6.243 | 1 | 0.012 | * |
| Genotype | 4.190 | 1 | 0.041 | * |
| Sex | 0.227 | 1 | 0.634 |  |
| Train | 454.062 | 1 | <2x10-16 | *** |
| Genotype x Sex | 3.048 | 1 | 0.081 | . |

